# A model of antigen processing improves prediction of MHC I-presented peptides

**DOI:** 10.1101/2020.03.28.013714

**Authors:** Timothy O’Donnell, Alex Rubinsteyn, Uri Laserson

## Abstract

Computational prediction of the peptides presented on MHC class I proteins is an important tool for studying T cell immunity. The data available to develop such predictors has expanded with the use of mass spec to identify naturally-presented MHC ligands. In addition to elucidating binding motifs, the identified ligands also reflect the antigen processing steps that occur prior to MHC binding. Here, we developed an integrated predictor of MHC I presentation that combines new models for MHC I binding and antigen processing. Considering only peptides first predicted by the binding model to bind strongly to MHC, the antigen processing model is trained to discriminate published mass spec-identified MHC I ligands from unobserved peptides. The integrated model outperformed the two individual components as well as NetMHCpan 4.0 and MixMHCpred 2.0.2 on held-out mass spec experiments. Our predictors are implemented in the MHCflurry package, version 1.6.0 (github.com/openvax/mhcflurry).

## Introduction

Cytotoxic T cells recognize peptides presented in complex with major histocompatibility class I (MHC I) molecules on cell surfaces. These peptides are usually derived from the degradation of endogenous proteins and comprise a snapshot of the protein content of the cell, enabling T cells to distinguish healthy cells from those with viral, bacterial, or tumor-associated mutated proteins^1,2^. The repertoire of MHC I-presented peptides is generated through a complex series of biochemical processes, beginning with cleavage of a protein into peptides in the proteosome, further cleavage (or destruction) by cytosolic peptidases, peptide transport into the endoplasmic reticulum (ER) through the transporter associated with antigen processing (TAP) complex, trimming by ER-resident aminopeptidases (ERAP), and stable association with one of the several MHC I proteins expressed by a cell^3^. The MHC I genes (*HLA-A, HLA-B*, and *HLA-C* in humans) are the sites of dramatic allelic variation at the population level, with each of the thousands of known *HLA* alleles associated with a strict, potentially distinct peptide binding preference. While a high-affinity interaction with MHC I is the most selective requirement for a peptide to be presented, the other processes in the antigen presentation pathway likely exert important secondary effects.

Prediction of MHC I-presented peptides is a critical tool in vaccine design and studies of infectious disease, autoimmunity, and cancer^4^. Most predictive pipelines in use today focus only on MHC I binding affinity (BA) prediction. While predictors fit to small datasets for individual antigen processing steps have been proposed,^5–8^ and integrated with binding affinity predictions to give a composite score^9–11^, improvements in accuracy from these approaches have been modest at best^12^. The relatively recent accumulation of large datasets of mass spec (MS)-identified MHC I ligands provides an opportunity revisit antigen processing prediction using larger and potentially more biologically-relevant datasets. While the antigen processing information in these datasets likely already informs existing MHC I binding predictors trained on MS datasets ^13,14^, antigen processing remains intertwined with MHC I binding preferences in these predictors, making it difficult to interpret the individual contributions and potentially leading to lower predictive accuracy.

In this work, we develop separate predictors for MHC allele-dependent effects (MHC I binding affinity; “BA”) and allele-independent effects (antigen processing; “AP”). We first trained a new pan-allele MHC I binding affinity predictor (referred to as MHCflurry 1.6.0 BA) on available MHC I ligand data, including affinity measurements and MS datasets. The use of *in vitro* affinity measurements in the training data, which are largely independent of antigen processing, is one of several design choices intended to limit the BA predictor’s tendency to learn antigen processing signals (Supplemental Note 1). We use the BA predictor to generate a training set for a model of antigen processing by combining MS-identified peptides (hits) with unobserved peptides (decoys), where both hits and decoys are predicted by the BA predictor to bind the relevant MHC I alleles. The antigen processing predictor thus models the residual allele-independent sequence properties that were not learned by the BA predictor. In support of its biological relevance, the processing predictor favored peptides consistent with established motifs for key antigen processing steps and showed quantitative agreement with an independent dataset of proteasome-cleaved peptides^15^. We combined the BA and AP models in a logistic regression model, which we refer to as the presentation score (MHCflurry 1.6.0 PS). Using a benchmark of held-out MS datasets, we found that the PS predictor outperformed both its component models and the commonly-used NetMHCpan 4.0 and MixMHCpred 2.0.2 predictors. The margin of improvement was substantial, with at least a 40% increase in positive predictive value (PPV) for all comparisons.

## Methods

### MS benchmark construction and approach

#### Dataset curation

To benchmark the binding predictors developed here and elsewhere, we collected datasets from 11 studies that identified MHC I-bound peptides using MS. We included only samples with known four-digit MHC I genotypes. Two of the studies included experiments that used cell lines engineered to express a single MHC I allele. We refer to these as the MONOALLELIC samples, which comprise 72 samples from the recent publication by Sarkizova et al^16^ and 8 samples from Abelin et al^17^. We refer to the other samples, in which exact MHC I restrictions were not experimentally determined, as the MULTIALLELIC samples. We divided these into two groups: MUTLIALLELIC-OLD, comprised of 56 experiments from eight studies published through 2018^18–25^, and MULTIALLELIC-RECENT, comprised of 20 experiments from two studies published in 2019^16,26^. Curated samples are listed in Table S1, the sample groups used for each benchmark experiment are given in Table S2, and Additional Files 1 and 2 give the full datasets for the MULTIALLELIC and MONOALLELIC benchmarks, respectively.

#### Decoy selection

We generated accuracy benchmarks from the curated datasets by sampling unobserved peptides (decoys) from the same proteins as the observed peptides (hits) for each sample. We first associated each sample with a publicly-available RNA-seq expression dataset based on its cell type or tissue of origin. Each MS-identified peptide was searched in the Uniprot human reference proteome (UP000005640_9606). In the case of multiple matches, we used the RNA-seq for each sample to select a single protein with the highest mRNA expression (total across transcripts). For each sample, we randomly selected *99n* decoy peptides, where *n* is the number of hits. Equal numbers of decoy peptides of each length (8, 9, 10, 11) were sampled. We excluded from this procedure all peptides (hits and decoys) that were present in the training data for any of a sample’s alleles for the MHCflurry predictor under evaluation, as well as hits that contained noncanonical amino acids or could not be matched to a Uniprot protein and Ensembl gene (Ensembl release 98).

#### Comparison to existing tools

We included two existing tools in our benchmarks, NetMHCpan 4.0 (ref. ^13^) and MixMHCpred 2.0.2 (ref. ^14,18^). The NetMHCpan 4.0 tool is an ensemble of neural networks trained on peptide-MHC I affinity measurements plus peptides from monoallelic MS experiments. It gives separate predictions for each kind of data, i.e. a BA prediction and an eluted ligand (EL) prediction, which we evaluated as separate predictors in our benchmarks. MixMHCpred is trained on peptides identified in multiallelic MS experiments, which it deconvolves into clusters that are subsequently associated with MHC I alleles. As both the MHCflurry binding affinity predictor and NetMHCpan 4.0 do not incorporate multiallelic MS in their training sets, we evaluated them on the full MULTIALLELIC benchmark. As MixMHCpred is trained on multiallelic MS, we tested it only on samples published recently (after it was trained), i.e. the MULTIALLELIC-RECENT benchmark. The two studies in the MONOALLELIC dataset were published after NetMHCpan 4.0 and MixMHCpred 2.0.2 were released, so it is also an appropriate test set for them. The released MHCflurry 1.6.0 binding predictor incorporates the MONOALLELIC dataset as training data, however, so in evaluations of this benchmark we used a variant of MHCflurry re-trained without these datasets. As the logistic regression model used in the MHCflurry 1.6.0 PS predictor was fit to the MULTIALLELIC-OLD subset, we benchmarked it only on the MULTIALLELIC-RECENT samples.

#### Scoring metrics

We assessed accuracy at distinguishing MS hits from decoys using two metrics, area under the curve (AUC) and positive predictive value (PPV). The AUC is a standard accuracy metric for classification tasks, interpretable as the probability that a randomly selected hit will be scored higher by a predictor than a randomly selected decoy. The PPV, as defined here, focuses on a predictor’s ability to rank hits far above the decoys. To calculate PPV, for each sample we sorted the *n* hits and *99n* decoys by their predictions, and calculated the fraction of the top *n* peptides that are hits. A random predictor would score 0.01 in PPV and 0.5 in AUC.

### MHC I binding affinity predictor

The MHCflurry 1.6.0 BA predictor is a new pan-allele MHC I binding affinity predictor that supports variable-length peptides up to 15-mers. It extends an earlier version of MHCflurry^27^ —in which separate neural networks were trained for each of 112 supported alleles— to now support 14,993 MHC I alleles using a single neural network ensemble. The input to each neural network consists of (1) an encoding of the peptide amino acid sequence and (2) an encoding of the amino acids at 37 selected positions from a multiple sequence alignment of a large number of MHC I alleles across species. The neural networks each output a nanomolar binding affinity (transformed to a 0-1 scale using the formula 1 − *log* _50000_ (*nM affinity*)). The mean of the transformed predictions over the ensemble gives the overall prediction. The predictor is trained on affinity measurements and MS datasets from cell lines monoallelic for a single MHC I allele. MS hits are assigned a “<50 nM” binding affinity. The models are trained using a variant of the mean squared loss (MSE) that supports such inequalities, as described previously^27^.

#### Peptide representation

Peptides of 15 amino acids or shorter are supported. Each peptide is transformed to a length-45 sequence by concatenating three representations: left aligned, centered, and right aligned. Unused positions are represented using a special X symbol, treated as a 21st amino acid. For example, the peptide “GILGFVFTL” is represented by concatenating “GILGFVFTLXXXXXX” (left aligned), “XXXGILGFVFTLXXX” (centered), and “XXXXXXGILGFVFTL” (right aligned), resulting in “GILGFVFTLXXXXXXXXXGILGFVFTLXXXXXXXXXGILGFVFTL”. Each amino acid in this sequence is transformed to a 21-dimensional vector using the BLOSUM62 substitution matrix^28^, extended to include the X placeholder, which is assigned similarity 1 to itself and 0 to all amino acids.

#### Allele representation

MHC I alleles are represented to the neural network by the amino acids at 37 positions from a global multiple sequence alignment. This representation is referred to as a “pseudosequence” by the NetMHCpan authors. We use the 34 peptide-contacting positions included in the NetMHCpan 4.0 pseudosequence (re-derived using a new alignment), plus 3 additional positions. The additional positions were selected to differentiate several pairs of alleles (A*23:01/A*24:13, A*29:01/A*29:02, B*44:02/B*44:27, C*03:03/C*03:04) that shared identical 34-mer NetMHCpan pseudosequences. Similar to the peptide encoding, each amino acid was transformed to a 21-dimensional vector using the BLOSUM62 substitution matrix, where a placeholder X character represents global alignment positions with no residue (i.e. a deletion) for a particular MHC allele.

#### Allele multiple sequence alignment

Full-length *HLA-A, -B, -C, -E, -F*, and *-G* amino acid sequences were downloaded from the IMGT/HLA^29^ project database, mouse H-2 sequences from uniprot^30^ (H-2 Db from accession P01899; H-2 Dd, P01900; H-2 Dp, P14427; H-2 Dk, P14426; H-2 Dq, Q31145; H-2 Kb, P01901; H-2 Kd, P01902; H-2 Kk, P04223; H-2 Kq, P14428; H-2 Ld, P01897; H-2 Lq, Q31151), and MHC I sequences for additional species from the IPD-MHC database^31^. After filtering to MHC I alleles with names parseable by the mhcnames package (https://github.com/openvax/mhcnames), the sequences were aligned using Clustal Omega 1.2.1 (ref. ^32^) with the command “clustalo -i class1.fasta -o class1.aligned.fasta.” Positions from the resulting alignment were selected that best recapitulated the NetMHCpan pseudosequences^13^, then extended to fully differentiate the 171 alleles with at least 50 entries in the training dataset. The final sequences included with MHCflurry encompass 14,993 MHC I alleles.

#### Training data

MHC ligand entries were downloaded from the Immune Epitope Database (IEDB)^33^ on Dec. 20, 2019. Entries with parseable MHC I allele names and peptides of length 8-15 with no post-translational modifications were retained. The MS data from IEDB was extended to include 47,459 additional MS hits from the SysteMHC Atlas project^34^ (downloaded on May 6, 2019) plus hits from the MONOALLELIC benchmark: 138,283 hits from Sarakizova et al16 and 4,808 hits from Abelin et al.17 The training set for the BA predictor is available as Additional File 3. For testing on the MONOALLELIC benchmark, a predictor variant trained without these datasets was used (Additional File 4). The set of IEDB affinity measurements was augmented with the “BD2013” dataset from Kim et al.^35^. The final training set consisted of 470,586 MS hits and 219,357 affinity measurements. A summary of the training datasets used for all predictors introduced in this work is given in Table S3.

#### Neural network architectures

Each neural network in the ensemble corresponds to one of 35 possible model architecture variants. The overall design in all cases is similar: the allele and peptide representations are concatenated, flattened, and passed through a series of two or three dense layers, whose size ranges from 256 to 1024. Each dense layer is followed by a Dropout layer^36^ with the dropout rate set to 50%. Architectures vary in terms of the size and number of the dense layers, the amount of regularization applied to dense layer weights (L1 penalty), and whether skip connections are used to give each dense layer direct access to the two preceding layers.

#### Model training

The training data was sampled four times to generate four training subsets. Each subset excluded one quarter or 100 of the training points for each MHC I allele, whichever was less for each allele. Models corresponding to the 35 architectures were fit to each of the four training subsets, for 140 trained models in total. Initial weights were selected using layer-sequential unit-variance initialization^37^. This was followed by a pre-training step, in which the network was fit to synthetic measurements generated by a previous version of MHCflurry (version 1.2.0, using allele-specific models trained without MS datasets). The synthetic data consists of affinity predictions for random peptides across 99 alleles. While pre-training on synthetic data, the training data was used as a test set for early stopping, i.e. pre-training was halted once the mean square error (MSE) was no longer improving on the actual (non-synthetic) training data. Typically, several hundred million synthetic measurements were used. Model fitting then proceeded using the training data, keeping 10% held-out for early stopping. The training dataset was augmented to include random peptides set to have a very weak affinity (>35,000 nM). The lengths of the random peptides were selected to equalize the number of non-binder data points across peptide lengths for each allele. As in NetMHC, the sequences of the random negative peptides were resampled after each epoch.

#### Model selection

From the 140 trained models, the model selection procedure selected nine to use in the final predictor ensemble. This was done by selecting a set of models from each of the four training subsets independently, using MSE on the held-out points as the accuracy metric. A forward stepwise procedure was used, in which models are added until the ensemble accuracy no longer improves. The final ensemble was the union of the models selected across training subsets.

### Antigen processing predictor

The MHCflurry 1.6.0 AP predictor is trained to model the MHC allele-independent effects that were not captured by the BA predictor. We designed it with proteasomal cleavage prediction in mind, although the resulting models are expected to capture a range of effects. Since the efficiency of cleavage by proteasomes and peptidases are affected by the residues both before and after the cut site, we experimented with the inclusion of the flanking sequence on either side of the peptide from its source protein. To understand the importance of flanking sequences, we trained two variants of the MHCflurry 1.6.0 AP predictor: one takes as input a peptide plus the 15 amino acids on either side in its source protein (AP with flanks) and one takes only a peptide (AP without flanks). Both models are independent of MHC allele.

#### Training data

The AP predictor is trained on the MS hits from the MONOALLELIC benchmark samples (Additional File 5). Hits of length 8-11 were matched 1:100 to decoys of the same length and from the same proteins. The resulting peptides for each sample were sorted by MHCflurry 1.6.0 BA prediction for the relevant allele, and the top 2% strongest binding peptides (hits and decoys) were selected for inclusion in the AP predictor’s training set. This resulted in a training set of 297,313 distinct peptides (399,392 entries total including duplicate peptides selected from different MS samples), of which 106,738 (36%) were hits. The median BA prediction for the peptides in AP training set was 44 nM for decoys and 23 nM for hits.

#### Neural network architectures

AP models were trained using 128 neural network architectures with a similar overall design but varying in layer sizes, activation function, level of L1 weight regularization, and dropout rate. The input to each network consists of peptides and flanking sequences (if used) as they occur in the source protein (N-flank - peptide - C-flank), with each amino acid encoded as a 21-dimensional vector using the BLOSUM62 substitution matrix extended to include an X symbol, as in the BA predictor. The sequence is right-padded with X characters to generate fixed-length inputs of length 45 (AP with flanks) or 15 (AP without flanks). The first layer is convolutional, with a kernel size of 11-17 and 256 or 512 filters, tanh or relu activation, and dropout. This transforms the input sequence into a new sequence of the same length but with up to 512 channels instead of 21. From this representation, two parallel convolutional layers with a kernel size of 1 (i.e. each position is considered independently) are applied to predict the favorability of an “N-terminal cut” and a “C-terminal cut” at each position in the sequence. The cut site predictors are implemented using two stacked convolutional layers with a kernel size of 1, and thus can be thought of as 2-layer dense networks that consider a single position in the learned representation of the input sequence. For example, in one model architecture the “C-terminal cut” predictor takes a 512-vector corresponding to a position in the input sequence and applies a dense layer with 8 outputs followed by a single-output dense layer. The final layer in the AP architecture is a dense layer that integrates these cut-site predictor results. It takes as input: (1) the N-terminal cut site prediction at the peptide N-terminus, (2) the max of the N-terminal cut site predictions across the rest of the peptide, (3) the C-terminal cut site prediction at the peptide C-terminus, and (4) the max of the C-terminal cut site predictions across the rest of the peptide. This design is motivated by the intuition that presented MHC I ligands must have favorable cleavability at their termini but avoid cleavage at interior residues. To give the model further opportunity to consider average properties of the flanking sequences (e.g. secondary structure), two additional inputs are given to the final layer: (5) the result from a dense layer applied to the per-channel average of the initial convolutional layer across the N-flank, (6) a similar result for the C terminus. The final layer in the AP predictor is thus intended to model the tradeoff between cleavability at the peptide termini, cleavability at interior positions in the peptide, and overall favorability of cleavage given the flanking sequences.

#### Model training and selection

The training data was sampled to generate four subsets, where each training subset held out 10 randomly selected MS experiments. Models corresponding to the 128 neural network architectural variants were fit to each training subset (512 models total). Models were trained using the Adam optimizer^38^ and binary cross entropy loss with a random 10% of each training subset used for early stopping. Model selection was performed for each training subset separately using AUC on the held-out data as the accuracy metric. The final selected ensemble included 8 models.

#### Evaluation on “mass spectrometry analysis of proteolytic peptides” (MAPP) data

Proteasome-cleaved peptides of lengths 8-11 were extracted from Wolf-Levy et al., Supplementary Data 2^15^. Peptides identified in any experiment (untreated or TNF/IFN-treated) were included. For each observed peptide (hit), a length-matched unobserved decoy peptide was randomly sampled from the same gene.

### Presentation score model

The MHCflurry 1.6.0 PS predictor is a two-input logistic regression model that integrates a BA prediction (tightest predicted binding affinity over the MHC I alleles for a sample) with an AP prediction to give a composite prediction, referred to as the presentation score. It has just three learned parameters: two coefficients and an intercept. In contrast to the BA and AP predictors, which use only monoallelic MS, the PS model is fit to multiallelic MS datasets. Training data for the PS model was generated using the MULTIALLELIC-OLD set of samples by sampling length-matched decoys (two decoys per hit) from the same proteins as the hits (Additional File 6). The training set included 753,777 entries (of which 251,277 were hits) from 56 samples. The model was fit using the logistic regression implementation in scikit-learn^39^ with the LBFGS solver and default parameters. Two variants of the PS model were generated: one that made use of the AP with flanks predictor (i.e. upstream and downstream amino acids included) and another that used the AP without flanks predictor. The PS model was benchmarked on the MULTIALLELIC-RECENT samples.

## Results

We first compared the new MHCflurry 1.6.0 binding affinity (BA) predictor (Fig. 1a) to existing binding affinity (NetMHCpan 4.0 BA) and MS ligand (NetMHCpan 4.0 EL and MixMHCpred 2.0.2) predictors on the MULTIALLELIC benchmark. MHCflurry BA performed best at differentiating MS hits from decoys in terms of AUC, outperforming NetMHCpan BA on 74 of 76 samples, NetMHCpan EL on 56 of 76, and MixMHCpred on 18 of 20 samples in the MULTIALLELIC-RECENT subset (Fig. 1b). The increase in AUC relative to NetMHCpan BA was 5.7% (bootstrap 95% confidence interval 4.7–6.7), 1.8% (1.3–2.5) relative to NetMHCpan EL, and 2.6% (1.8–3.4) relative to MixMHCpred, with the greatest improvements observed for non-9-mer peptides (Fig. 1c). By contrast, in terms of PPV, which emphasizes differences at the high end of a predictor’s output, MHCflurry BA showed only an advantage in comparison to NetMHCpan BA, with a mean improvement in PPV of 50% (39 - 65). Differences were within error in comparison to the MS ligand predictors, with a mean 3.7% (−2.0 - +10.3) improvement over NetMHCpan EL and a trend toward underperforming MixMHCpred, with a mean difference of -5.0% (−12.6 - +4.6; Fig. 1d). Similar results were observed when evaluating the predictors on the MONOALLELIC benchmark using a variant of MHCflurry that was trained without the MONOALLELIC samples (Fig. S1).

**Figure 1.**
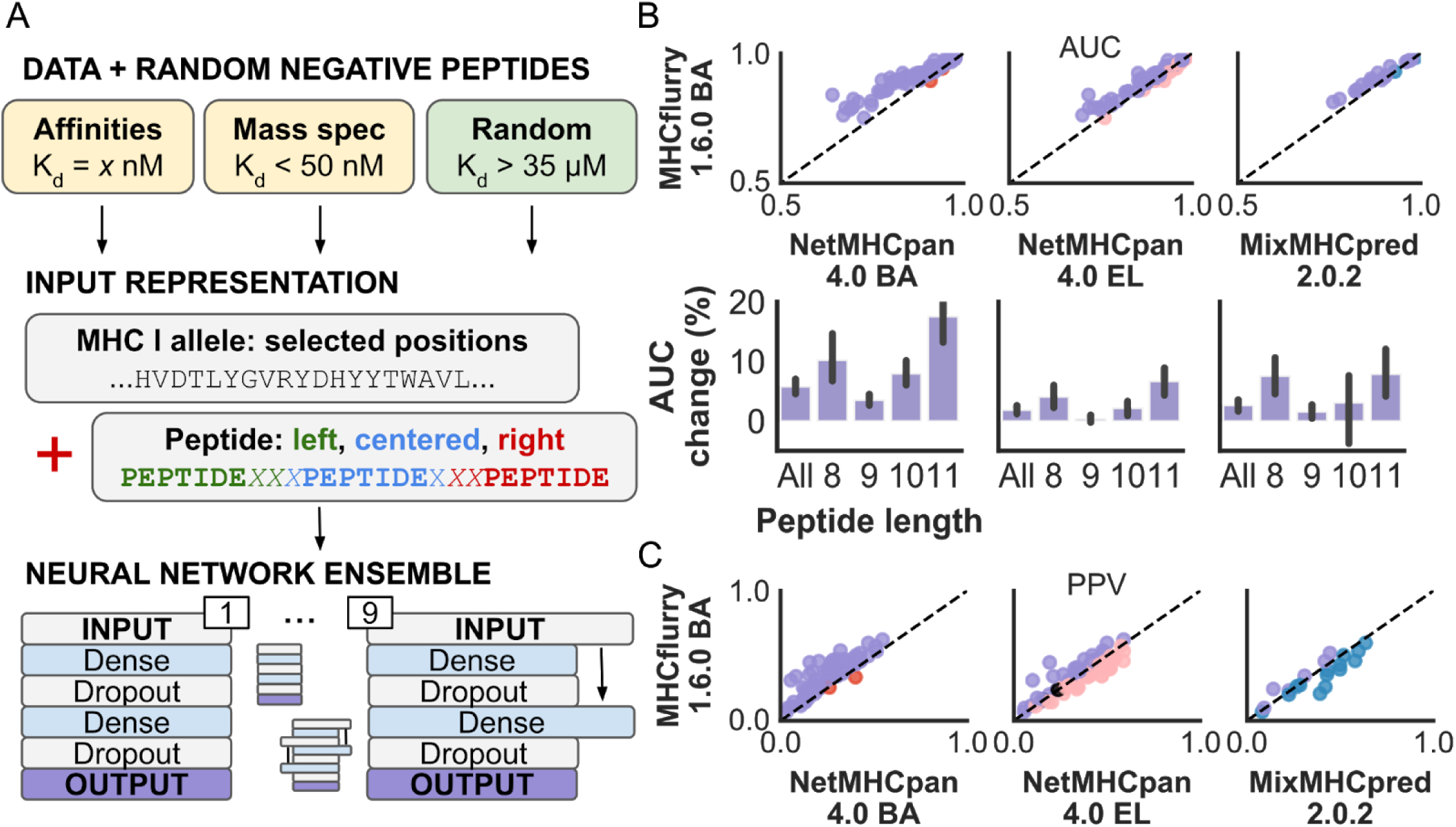
MHCflurry 1.6.0 binding affinity (BA) predictor architecture and benchmark. **(a)** BA predictor training data, model input representations, and neural network architectures. **(b)** Area under the curve (AUC) of MHCflurry 1.6.0 BA in comparison to other predictors across the MULTIALLELIC benchmark experiments. Each point in the top panel corresponds to a single experiment from the MULTIALLELIC (NetMHCpan 4.0 BA and EL) or MULTIALLELIC-RECENT (MixMHCpred 2.0.2) benchmarks. The bottom panel indicates mean improvement in AUC across experiments per peptide length. Error bars are bootstrap 95% confidence intervals of the mean. **(c)** Positive predictive value (PPV) of MHCflurry 1.6.0 BA in comparison to other predictors. Points correspond to the same experiments as in (a).

We hypothesized that explicitly modeling processes that do not depend on MHC I allele might enable accuracy improvements over MHCflurry BA alone. We therefore developed an MHC I allele-independent model trained to distinguish hits from decoys where both the hits and decoy peptides are predicted to be tight binders (rank less than 2%) by the MHCflurry BA predictor. We refer to this model as the MHCflurry 1.6.0 antigen processing (AP) predictor. Its neural network architecture is motivated by the possibility of learning peptide N- and C-terminal cleavage or processing signals (Fig. 2a). We trained two versions of the AP predictor on the MONOALLELIC benchmark dataset. One predictor includes the peptide plus the 15 immediately upstream and downstream residues from its source protein (AP with flanks); the second predictor includes only the peptide (AP without flanks).

**Figure 2.**
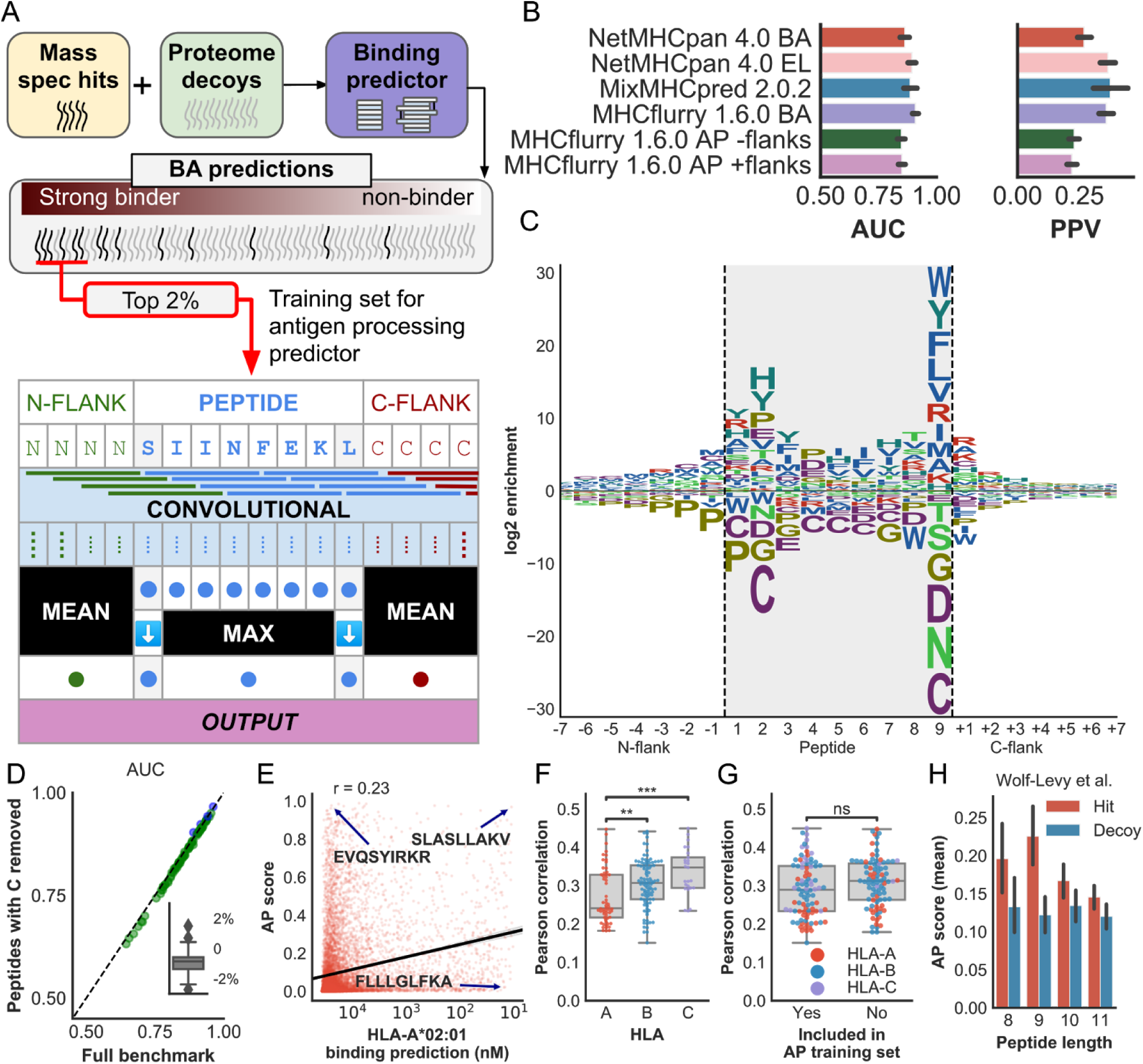
The MHCflurry 1.6.0 antigen processing (AP) predictor models MHC I allele-independent effects. **(a)** AP predictor training scheme and neural network architecture. **(b)** Mean AUC and PPV accuracy on the multiallelic mass spec benchmarks for the AP predictor in comparison to other predictors. The MixMHCpred 2.0.2 tool was benchmarked on the MULTIALLELIC-RECENT subset; the other predictors were tested on the full MULTIALLELIC benchmark. **(c)** Sequence logo for the motif learned by the AP predictor. Positive values (above the center line) indicate enrichments above proteome level; negative values indicate depletions. **(d)** Comparison of AP predictor AUC scores on the MULTIALLELIC benchmark samples when peptides containing cysteine were removed (y-axis) vs. retained (x-axis). The inset shows the percentage change in AUC when cysteine-containing peptides are removed. **(e)** Correlation of the AP predictor with the BA prediction for HLA-A*02:01. Each red dot corresponds to a 9-mer peptide sampled from the proteome. The black line indicates the best-fit regression line, and example peptides are indicated. **(f, g)** Correlation of AP predictor with the BA prediction across HLA alleles by gene (f) or by allele representation in the AP training set (g). **(h)** Mean AP score for proteasome-associated peptides observed by Wolf-Levy et al.15 (hits) and unobserved peptides from the same genes (decoys).

To understand if the MHCflurry AP predictors learned a meaningful signal, we evaluated their accuracy on the MULTIALLELIC benchmark (Fig. 2b). While the AP variants underperformed the standard MHC binding predictors and MHCflurry BA, they performed better than might be expected given that they do not take MHC allele as an input. Both the AP with flanks and AP without flanks predictors showed a mean 0.85 AUC (bootstrap 95% CI 0.83 - 0.87), compared to 0.91 (0.90 - 0.92) for the MHCflurry BA predictor (Fig. 2b). This suggested that the MHCflurry AP predictors had learned a meaningful MHC I allele-independent signal from the MONOALLELIC MS training set.

To understand what the MHCflurry AP predictors may be learning, we ranked all 9-mer peptides in the MULTIALLELIC benchmark by AP prediction, calculated position weight matrices for the top 1% highest predictions, and plotted a sequence logo (Fig. 2c and Additional File 7). This analysis showed the AP predictor learned that hits are depleted for cysteines across the peptide, a known bias of MS^40^. It also showed depletion of prolines from the first position in the peptide (P1) extending upstream into the N-flank, as well as strong but complex specificity at the C-terminus of the peptide and some preferences at the downstream flanking residue. These signals suggested qualitative agreement with established signatures for TAP transport, proteasomal cleavage, and ERAP trimming (see Discussion).

As cysteine depletion is one of the strongest known MS biases, we were concerned that much of the accuracy of the AP predictor may be due its modeling of this bias. We therefore repeated the AUC analysis on the MULTIALLELIC benchmark after removing peptides (hits and decoys) that contained cysteine. This analysis showed very similar AUC values as the earlier analysis, with a change in AUC of less than 3% for all samples (Fig. 2d). Thus, while the AP predictor learns the cysteine MS bias, this effect alone is not the primary driver of its performance.

To evaluate the extent to which the AP predictors learned a signal that is also learned by the MHCflurry BA predictor, we measured the correlation between AP and BA predictions for random peptides across the *HLA-A, HLA-B*, and *HLA-C* alleles in the BA training set (n=181). The correlations were positive, significant, but modest in magnitude (Pearson r < 0.5) for all alleles tested. For example, AP and BA predictions for the HLA-A*02:01 allele showed a Pearson correlation of 0.23. While peptides predicted to bind HLA-A*02:01 tightly tended to have higher AP scores, it was possible to find peptides with high scores for one predictor but not the other (Fig. 2e). Correlations with the AP predictor were somewhat higher for *HLA-B* and *HLA-C* than for *HLA-A* alleles (Fig. 2f). The alleles used to train the AP predictor showed no greater AP vs. BA correlations on average than those that were not included in the AP training set (Fig. 2g). Overall, this analysis suggested that the AP predictor is at least partially non-redundant with the MHCflurry BA predictor.

To quantitatively assess if the AP predictor captures biologically-relevant effects, we tested it on an independent dataset of proteosome-cleaved peptides. We applied the AP predictor to 3,017 peptides identified by Wolf-Levy et al. using the “mass spectrometry analysis of proteolytic peptides” (MAPP) assay, in which cleaved peptides are reversibly cross-linked to cellular proteosomes and identified by MS^15^. AP predictor scores were significantly higher for MAPP-identified peptides (hits) than unobserved (decoy) peptides drawn from the same genes (Mann-Whitney p < 0.01 for each peptide length 8-11; Fig. 2h and Additional File 8). This indicated that the AP predictor learned a signal consistent with a key antigen processing step.

We next asked if the AP predictor may be combined with the MHCflurry BA predictor to achieve higher performance than either alone. We trained a logistic regression model that takes two inputs: the strongest MHCflurry BA prediction across alleles for the sample (transformed to fall in the range 0.0 - 1.0, with higher indicating a stronger binder), and the AP prediction (Fig. 3a). We refer to this logistic regression model as the MHCflurry presentation score (PS) predictor. We trained MHCflurry PS predictors using either the AP with flanking (PS with flanking) or AP without flanking (PS without flanking) predictors on the MULTIALLELIC-OLD dataset and evaluated performance on the MULTIALLELIC-RECENT benchmark.

**Figure 3.**
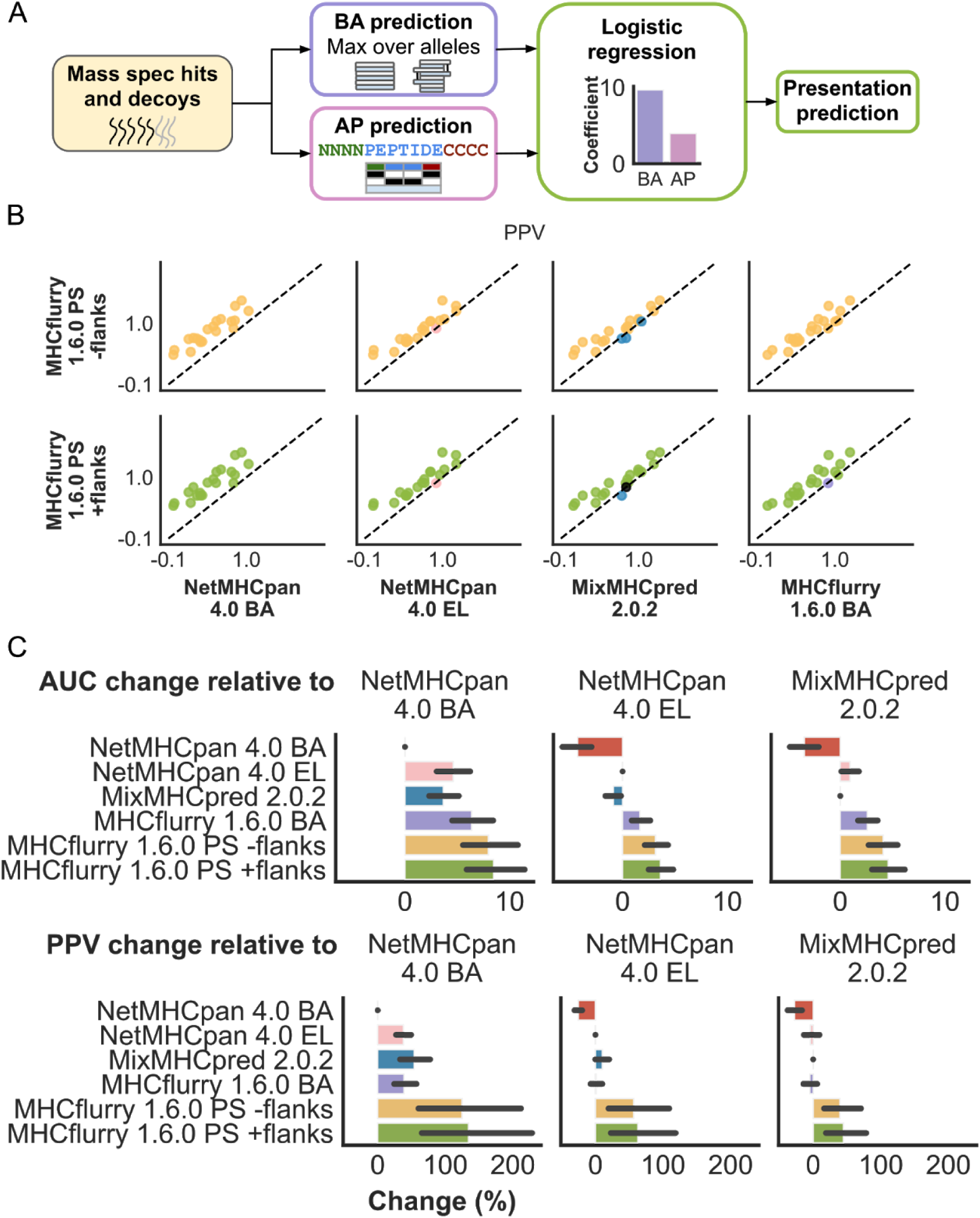
The MHCflurry 1.6.0 presentation score (PS) model combines binding affinity and antigen processing prediction. **(a)** The PS model is a two-input logistic regression model that integrates BA and AP predictions to give a composite score. It is trained on multiallelic mass spec hits and decoys. **(b)** Comparison of PPV scores of the PS models with other predictors. The top row shows the performance of the PS model variant that uses the AP without flanks predictor, which considers only the peptide and not the upstream and downstream flanking sequences. The bottom row corresponds to the variant that also considers the flanking sequences. **(c)** Mean percent change in AUC and PPV for the indicated predictors (y-axis) relative to each of the three existing predictors (columns). Error bars indicate 95% confidence intervals for the mean change.

Both PS models showed significantly improved accuracy, in terms of both AUC and PPV, over all other predictors tested (Fig. 3b,c). In terms of PPV, the PS without flanks predictor showed a 125% (64 - 190) improvement over NetMHCpan 4.0 BA, 57% (21 - 94) improvement over NetMHCpan 4.0 EL, 40% (19 - 64) improvement over MixMHCpred 2.0.2, and 49% (28 - 75) improvement over MHCflurry 1.6.0 BA. The use of flanking sequences made a small but consistent difference, further improving PPV by a mean 3.3% (1.8 - 4.8).

## Discussion

Our MHC I ligand prediction method is comprised of two neural network models: the MHCflurry BA predictor and the MHC I allele-independent MHCflurry AP predictor. The AP predictor is trained to learn what the BA predictor missed, i.e. residual sequence properties that distinguish hits from decoys among peptides predicted to bind MHC I tightly. Both predictors are trained on monoallelic MS datasets (plus affinity measurements for the BA predictor), and their results combined using a logistic regression model fit to multiallelic MS datasets. When evaluated on held-out multiallelic MS experiments, the combined predictor, referred to as MHCflurry presentation score (PS), outperformed the individual components and standard tools. While inclusion of flanking sequences, i.e. the adjacent residues in the peptide’s source protein, provided a small additional accuracy boost, the overall improvement over standard tools was also evident when only the peptide was provided to the AP predictor.

Our work builds on the approach by Abelin et al., who developed a proteasomal cleavage predictor fit to MS-identified MHC I ligands and observed an increase in accuracy when it was included along with other features in a logistic regression model^40^. Their work used a different method to control for MHC I binding signals in the cleavage predictor training set, however, in which decoys were selected to match the hit peptide amino acids at the first and last two positions, which encompass the anchor positions for most alleles. This is expected to disrupt the cleavage model’s ability to learn features at these positions, which are also the positions where we observed the strongest preferences.

The AP predictor sequence motif (Fig. 2c) shows similarities with the established preferences of key antigen processing steps. Deconvolution of effects, however, is complicated by the overlapping specificity of these steps, potentially a consequence of coevolution of the antigen processing machinery^7^. In particular, the AP predictor’s preferences at the C terminus of the peptide may reflect TAP binding and/or proteasomal cleavage. Work by Tampé and colleagues^41^ found that TAP favors peptides with a C terminal Phe, Tyr, Arg, or Leu and disfavors Asp, Glu, Asn and Ser, all recapitulated by the AP predictor. The AP motif is also consistent with the effects of the chymotryptic-like activity of the proteasome (cleavage after Phe, Tyr, Leu, Trp but not Gly) as well as tryptic-like (cleavage after Arg and Lys), but not caspase-like specificity^42,43^. Some agreement with proteasomal processing is also apparent at the interior residues of the peptide, such as a depletion of proline at P8 and enrichment for proline at P6 (referred to as P2 and P4, respectively, in cleavage studies). At the first flanking residue downstream of the peptide, which is free from the effects of TAP and MHC binding, the enrichments for Arg, Ala, Lys, Ser, and Gly are consistent with proteasomal cleavage preferences for the position after the cut site^15,42^, although the enrichment for Arg and Lys could also indicate tryptic-like specificity working from the C-terminus of the protein^40^. The striking depletion of prolines in the N-flanking residues up to and including the first position of the peptide (P1) and the enrichment for proline at P2 is consistent with trimming by ERAP^44^, although again we cannot exclude a contribution from proteasomal cleavage. These observations and the higher AP scores for proteasome-cleaved peptides identified by MAPP (Fig. 2h) suggest that the AP predictor has learned certain antigen processing signals, although a detailed deconvolution of effects remains future work.

An important limitation of this work is that we apply datasets of MHC I ligands detected by MS both to train and to benchmark our predictors. Assay biases, which we expect are modeled by the AP predictor, have the potential to erroneously inflate our accuracy scores. While the main known bias, depletion of cysteine, does not seem to have a dramatic effect on AP predictive accuracy, we cannot rule out the contributions of other kinds of bias. Our work also only addresses the steps contributing to MHC I ligand presentation, not T cell recognition of presented epitopes. Future work will need to assess whether the predictors described here enable improved prediction of T cell epitopes.

### Data and software availability

The predictors described here are available under an Apache 2 open source license in the MHCflurry package (https://github.com/openvax/mhcflurry). The datasets used to train and benchmark the predictors are deposited in Mendeley Data at https://data.mendeley.com/datasets/zx3kjzc3yx.

## Acknowledgements

This work was supported in part through the computational resources and staff expertise provided by Scientific Computing at the Icahn School of Medicine at Mount Sinai.

## Author contributions

T.O. designed research, performed research, and wrote the paper. A.R and U.L advised on data interpretation. All authors critically reviewed the manuscript.

## Supplemental figures and tables

**Table S1.**
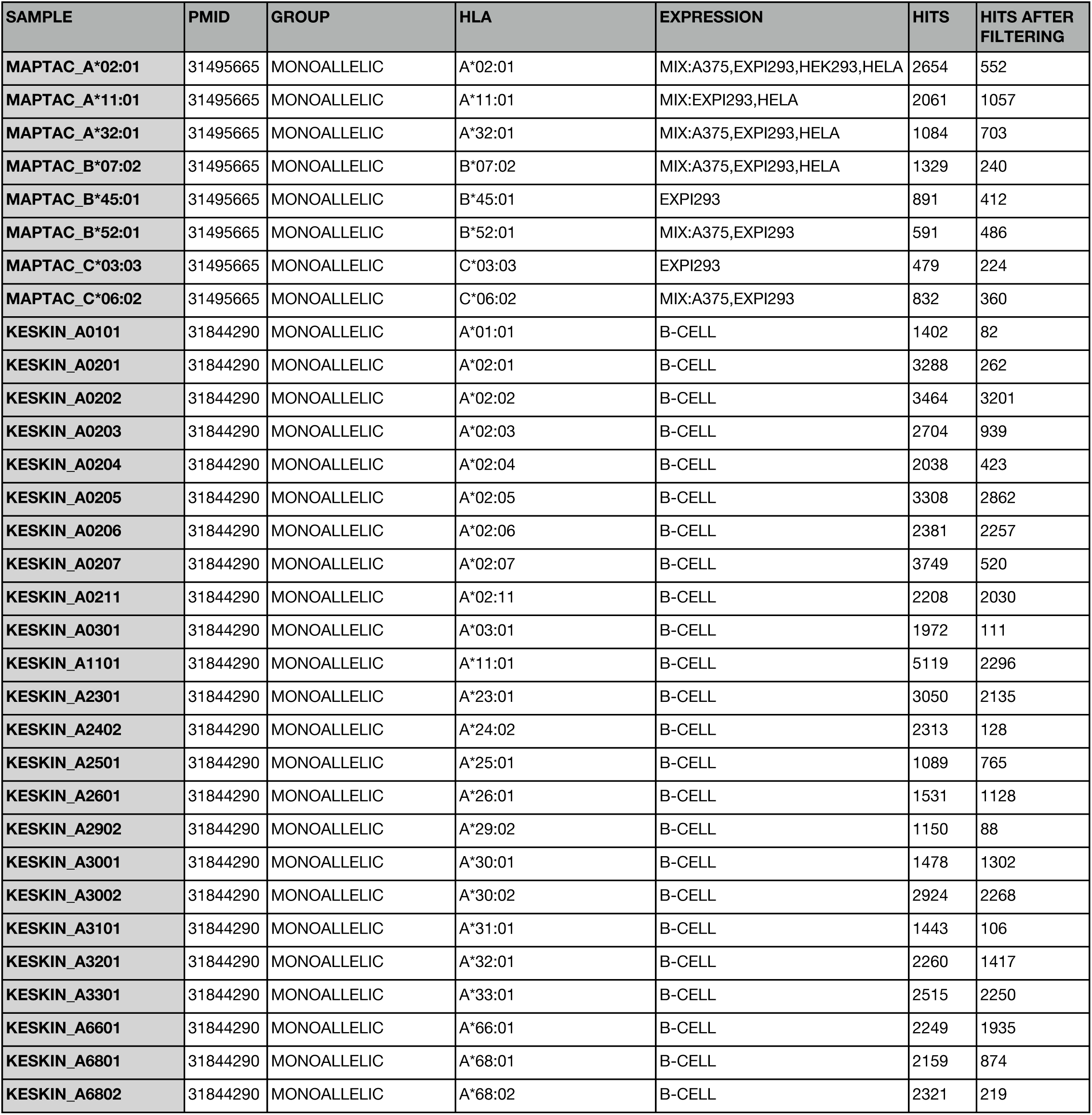
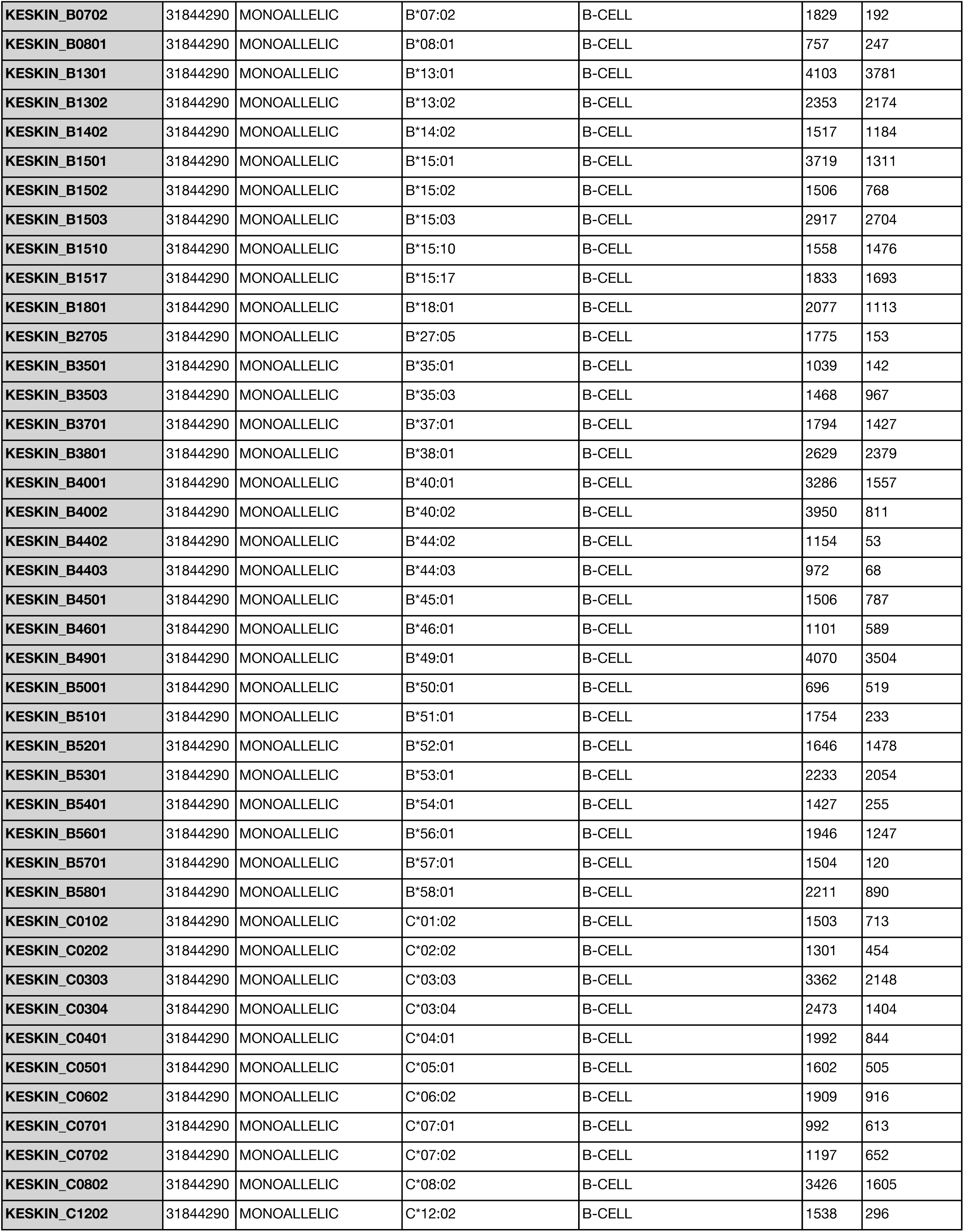
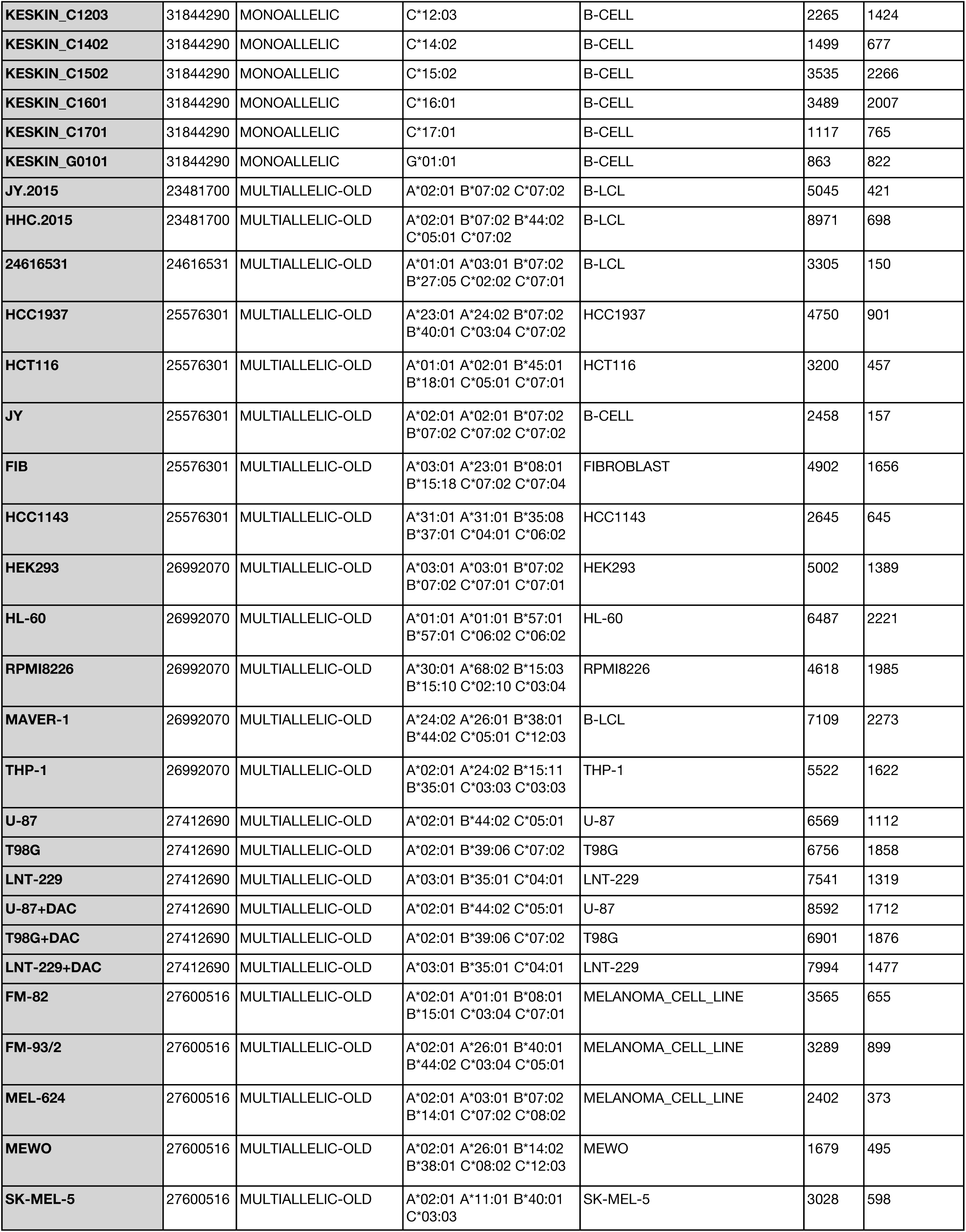
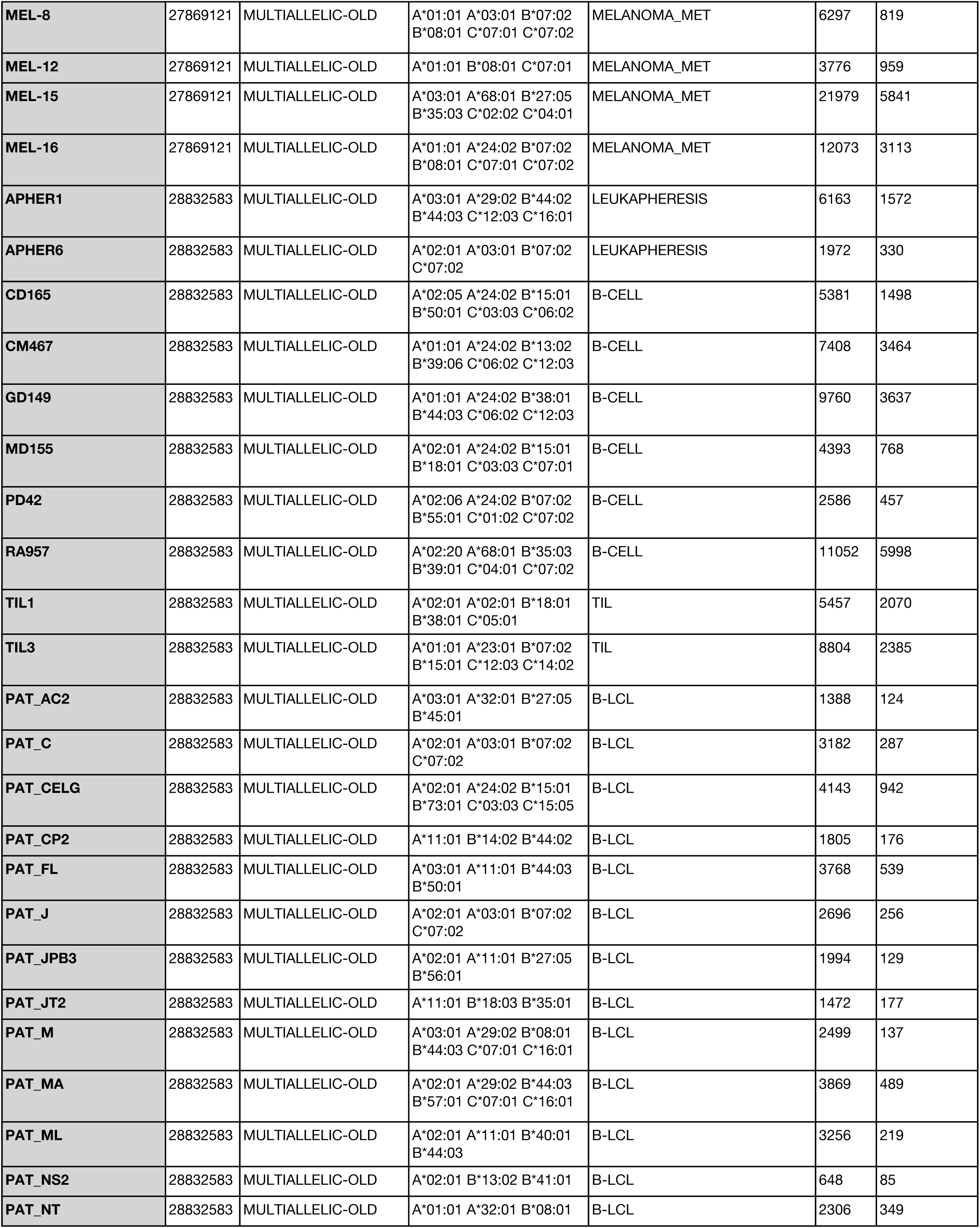
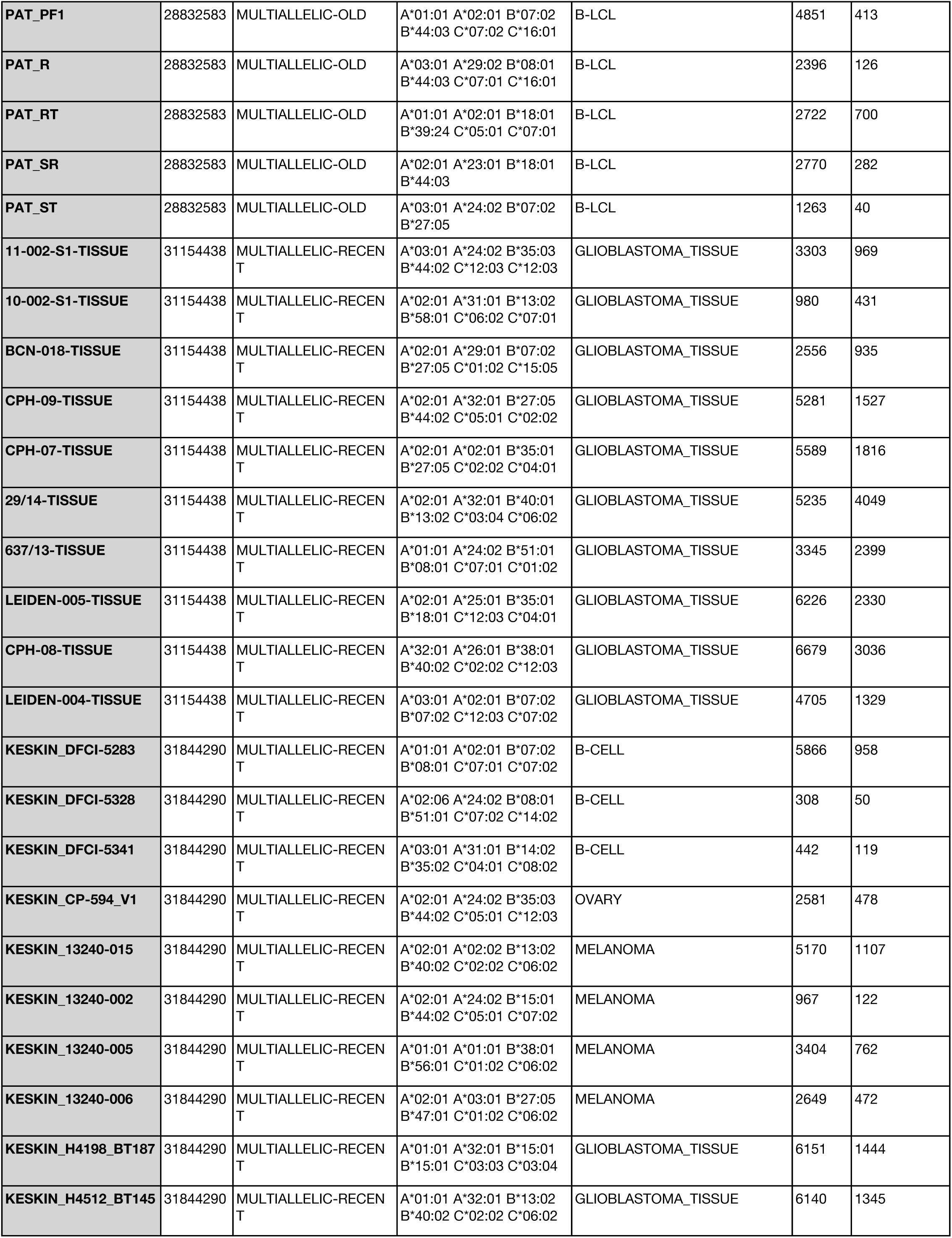
Published MHC I mass spec datasets used for benchmarking in this work.

| SAMPLE | PMID | GROUP | HLA | EXPRESSION | HITS | HITS AFTER FILTERING |
| --- | --- | --- | --- | --- | --- | --- |
| MAPTAC_A*02:01 | 31495665 | MONOALLELIC | A*02:01 | MIX:A375,EXPI293,HEK293,HELA | 2654 | 552 |
| MAPTAC_A*11:01 | 31495665 | MONOALLELIC | A*11:01 | MIX:EXPI293,HELA | 2061 | 1057 |
| MAPTAC_A*32:01 | 31495665 | MONOALLELIC | A*32:01 | MIX:A375,EXPI293,HELA | 1084 | 703 |
| MAPTAC_B*07:02 | 31495665 | MONOALLELIC | B*07:02 | MIX:A375,EXPI293,HELA | 1329 | 240 |
| MAPTAC_B*45:01 | 31495665 | MONOALLELIC | B*45:01 | EXPI293 | 891 | 412 |
| MAPTAC_B*52:01 | 31495665 | MONOALLELIC | B*52:01 | MIX:A375,EXPI293 | 591 | 486 |
| MAPTAC_C*03:03 | 31495665 | MONOALLELIC | C*03:03 | EXPI293 | 479 | 224 |
| MAPTAC_C*06:02 | 31495665 | MONOALLELIC | C*06:02 | MIX:A375,EXPI293 | 832 | 360 |
| KESKIN_A0101 | 31844290 | MONOALLELIC | A*01:01 | B-CELL | 1402 | 82 |
| KESKIN_A0201 | 31844290 | MONOALLELIC | A*02:01 | B-CELL | 3288 | 262 |
| KESKIN_A0202 | 31844290 | MONOALLELIC | A*02:02 | B-CELL | 3464 | 3201 |
| KESKIN_A0203 | 31844290 | MONOALLELIC | A*02:03 | B-CELL | 2704 | 939 |
| KESKIN_A0204 | 31844290 | MONOALLELIC | A*02:04 | B-CELL | 2038 | 423 |
| KESKIN_A0205 | 31844290 | MONOALLELIC | A*02:05 | B-CELL | 3308 | 2862 |
| KESKIN_A0206 | 31844290 | MONOALLELIC | A*02:06 | B-CELL | 2381 | 2257 |
| KESKIN_A0207 | 31844290 | MONOALLELIC | A*02:07 | B-CELL | 3749 | 520 |
| KESKIN_A0211 | 31844290 | MONOALLELIC | A*02:11 | B-CELL | 2208 | 2030 |
| KESKIN_A0301 | 31844290 | MONOALLELIC | A*03:01 | B-CELL | 1972 | 111 |
| KESKIN_A1101 | 31844290 | MONOALLELIC | A*11:01 | B-CELL | 5119 | 2296 |
| KESKIN_A2301 | 31844290 | MONOALLELIC | A*23:01 | B-CELL | 3050 | 2135 |
| KESKIN_A2402 | 31844290 | MONOALLELIC | A*24:02 | B-CELL | 2313 | 128 |
| KESKIN_A2501 | 31844290 | MONOALLELIC | A*25:01 | B-CELL | 1089 | 765 |
| KESKIN_A2601 | 31844290 | MONOALLELIC | A*26:01 | B-CELL | 1531 | 1128 |
| KESKIN_A2902 | 31844290 | MONOALLELIC | A*29:02 | B-CELL | 1150 | 88 |
| KESKIN_A3001 | 31844290 | MONOALLELIC | A*30:01 | B-CELL | 1478 | 1302 |
| KESKIN_A3002 | 31844290 | MONOALLELIC | A*30:02 | B-CELL | 2924 | 2268 |
| KESKIN_A3101 | 31844290 | MONOALLELIC | A*31:01 | B-CELL | 1443 | 106 |
| KESKIN_A3201 | 31844290 | MONOALLELIC | A*32:01 | B-CELL | 2260 | 1417 |
| KESKIN_A3301 | 31844290 | MONOALLELIC | A*33:01 | B-CELL | 2515 | 2250 |
| KESKIN_A6601 | 31844290 | MONOALLELIC | A*66:01 | B-CELL | 2249 | 1935 |
| KESKIN_A6801 | 31844290 | MONOALLELIC | A*68:01 | B-CELL | 2159 | 874 |
| KESKIN_A6802 | 31844290 | MONOALLELIC | A*68:02 | B-CELL | 2321 | 219 |
| KESKIN_B0702 | 31844290 | MONOALLELIC | B*07:02 | B-CELL | 1829 | 192 |
| KESKIN_B0801 | 31844290 | MONOALLELIC | B*08:01 | B-CELL | 757 | 247 |
| KESKIN_B1301 | 31844290 | MONOALLELIC | B*13:01 | B-CELL | 4103 | 3781 |
| KESKIN_B1302 | 31844290 | MONOALLELIC | B*13:02 | B-CELL | 2353 | 2174 |
| KESKIN_B1402 | 31844290 | MONOALLELIC | B*14:02 | B-CELL | 1517 | 1184 |
| KESKIN_B1501 | 31844290 | MONOALLELIC | B*15:01 | B-CELL | 3719 | 1311 |
| KESKIN_B1502 | 31844290 | MONOALLELIC | B*15:02 | B-CELL | 1506 | 768 |
| KESKIN_B1503 | 31844290 | MONOALLELIC | B*15:03 | B-CELL | 2917 | 2704 |
| KESKIN_B1510 | 31844290 | MONOALLELIC | B*15:10 | B-CELL | 1558 | 1476 |
| KESKIN_B1517 | 31844290 | MONOALLELIC | B*15:17 | B-CELL | 1833 | 1693 |
| KESKIN_B1801 | 31844290 | MONOALLELIC | B*18:01 | B-CELL | 2077 | 1113 |
| KESKIN_B2705 | 31844290 | MONOALLELIC | B*27:05 | B-CELL | 1775 | 153 |
| KESKIN_B3501 | 31844290 | MONOALLELIC | B*35:01 | B-CELL | 1039 | 142 |
| KESKIN_B3503 | 31844290 | MONOALLELIC | B*35:03 | B-CELL | 1468 | 967 |
| KESKIN_B3701 | 31844290 | MONOALLELIC | B*37:01 | B-CELL | 1794 | 1427 |
| KESKIN_B3801 | 31844290 | MONOALLELIC | B*38:01 | B-CELL | 2629 | 2379 |
| KESKIN_B4001 | 31844290 | MONOALLELIC | B*40:01 | B-CELL | 3286 | 1557 |
| KESKIN_B4002 | 31844290 | MONOALLELIC | B*40:02 | B-CELL | 3950 | 811 |
| KESKIN_B4402 | 31844290 | MONOALLELIC | B*44:02 | B-CELL | 1154 | 53 |
| KESKIN_B4403 | 31844290 | MONOALLELIC | B*44:03 | B-CELL | 972 | 68 |
| KESKIN_B4501 | 31844290 | MONOALLELIC | B*45:01 | B-CELL | 1506 | 787 |
| KESKIN_B4601 | 31844290 | MONOALLELIC | B*46:01 | B-CELL | 1101 | 589 |
| KESKIN_B4901 | 31844290 | MONOALLELIC | B*49:01 | B-CELL | 4070 | 3504 |
| KESKIN_B5001 | 31844290 | MONOALLELIC | B*50:01 | B-CELL | 696 | 519 |
| KESKIN_B5101 | 31844290 | MONOALLELIC | B*51:01 | B-CELL | 1754 | 233 |
| KESKIN_B5201 | 31844290 | MONOALLELIC | B*52:01 | B-CELL | 1646 | 1478 |
| KESKIN_B5301 | 31844290 | MONOALLELIC | B*53:01 | B-CELL | 2233 | 2054 |
| KESKIN_B5401 | 31844290 | MONOALLELIC | B*54:01 | B-CELL | 1427 | 255 |
| KESKIN_B5601 | 31844290 | MONOALLELIC | B*56:01 | B-CELL | 1946 | 1247 |
| KESKIN_B5701 | 31844290 | MONOALLELIC | B*57:01 | B-CELL | 1504 | 120 |
| KESKIN_B5801 | 31844290 | MONOALLELIC | B*58:01 | B-CELL | 2211 | 890 |
| KESKIN_C0102 | 31844290 | MONOALLELIC | C*01:02 | B-CELL | 1503 | 713 |
| KESKIN_C0202 | 31844290 | MONOALLELIC | C*02:02 | B-CELL | 1301 | 454 |
| KESKIN_C0303 | 31844290 | MONOALLELIC | C*03:03 | B-CELL | 3362 | 2148 |
| KESKIN_C0304 | 31844290 | MONOALLELIC | C*03:04 | B-CELL | 2473 | 1404 |
| KESKIN_C0401 | 31844290 | MONOALLELIC | C*04:01 | B-CELL | 1992 | 844 |
| KESKIN_C0501 | 31844290 | MONOALLELIC | C*05:01 | B-CELL | 1602 | 505 |
| KESKIN_C0602 | 31844290 | MONOALLELIC | C*06:02 | B-CELL | 1909 | 916 |
| KESKIN_C0701 | 31844290 | MONOALLELIC | C*07:01 | B-CELL | 992 | 613 |
| KESKIN_C0702 | 31844290 | MONOALLELIC | C*07:02 | B-CELL | 1197 | 652 |
| KESKIN_C0802 | 31844290 | MONOALLELIC | C*08:02 | B-CELL | 3426 | 1605 |
| KESKIN_C1202 | 31844290 | MONOALLELIC | C*12:02 | B-CELL | 1538 | 296 |
| KESKIN_C1203 | 31844290 | MONOALLELIC | C*12:03 | B-CELL | 2265 | 1424 |
| KESKIN_C1402 | 31844290 | MONOALLELIC | C*14:02 | B-CELL | 1499 | 677 |
| KESKIN_C1502 | 31844290 | MONOALLELIC | C*15:02 | B-CELL | 3535 | 2266 |
| KESKIN_C1601 | 31844290 | MONOALLELIC | C*16:01 | B-CELL | 3489 | 2007 |
| KESKIN_C1701 | 31844290 | MONOALLELIC | C*17:01 | B-CELL | 1117 | 765 |
| KESKIN_G0101 | 31844290 | MONOALLELIC | G*01:01 | B-CELL | 863 | 822 |
| JY.2015 | 23481700 | MULTIALLELIC-OLD | A*02:01 B*07:02 C*07:02 | B-LCL | 5045 | 421 |
| HHC.2015 | 23481700 | MULTIALLELIC-OLD | A*02:01 B*07:02 B*44:02<br>C*05:01 C*07:02 | B-LCL | 8971 | 698 |
| 24616531 | 24616531 | MULTIALLELIC-OLD | A*01:01 A*03:01 B*07:02<br>B*27:05 C*02:02 C*07:01 | B-LCL | 3305 | 150 |
| HCC1937 | 25576301 | MULTIALLELIC-OLD | A*23:01 A*24:02 B*07:02<br>B*40:01 C*03:04 C*07:02 | HCC1937 | 4750 | 901 |
| HCT116 | 25576301 | MULTIALLELIC-OLD | A*01:01 A*02:01 B*45:01<br>B*18:01 C*05:01 C*07:01 | HCT116 | 3200 | 457 |
| JY | 25576301 | MULTIALLELIC-OLD | A*02:01 A*02:01 B*07:02<br>B*07:02 C*07:02 C*07:02 | B-CELL | 2458 | 157 |
| FIB | 25576301 | MULTIALLELIC-OLD | A*03:01 A*23:01 B*08:01<br>B*15:18 C*07:02 C*07:04 | FIBROBLAST | 4902 | 1656 |
| HCC1143 | 25576301 | MULTIALLELIC-OLD | A*31:01 A*31:01 B*35:08<br>B*37:01 C*04:01 C*06:02 | HCC1143 | 2645 | 645 |
| HEK293 | 26992070 | MULTIALLELIC-OLD | A*03:01 A*03:01 B*07:02<br>B*07:02 C*07:01 C*07:01 | HEK293 | 5002 | 1389 |
| HL-60 | 26992070 | MULTIALLELIC-OLD | A*01:01 A*01:01 B*57:01<br>B*57:01 C*06:02 C*06:02 | HL-60 | 6487 | 2221 |
| RPMI8226 | 26992070 | MULTIALLELIC-OLD | A*30:01 A*68:02 B*15:03<br>B*15:10 C*02:10 C*03:04 | RPMI8226 | 4618 | 1985 |
| MAVER-1 | 26992070 | MULTIALLELIC-OLD | A*24:02 A*26:01 B*38:01<br>B*44:02 C*05:01 C*12:03 | B-LCL | 7109 | 2273 |
| THP-1 | 26992070 | MULTIALLELIC-OLD | A*02:01 A*24:02 B*15:11<br>B*35:01 C*03:03 C*03:03 | THP-1 | 5522 | 1622 |
| U-87 | 27412690 | MULTIALLELIC-OLD | A*02:01 B*44:02 C*05:01 | U-87 | 6569 | 1112 |
| T98G | 27412690 | MULTIALLELIC-OLD | A*02:01 B*39:06 C*07:02 | T98G | 6756 | 1858 |
| LNT-229 | 27412690 | MULTIALLELIC-OLD | A*03:01 B*35:01 C*04:01 | LNT-229 | 7541 | 1319 |
| U-87+DAC | 27412690 | MULTIALLELIC-OLD | A*02:01 B*44:02 C*05:01 | U-87 | 8592 | 1712 |
| T98G+DAC | 27412690 | MULTIALLELIC-OLD | A*02:01 B*39:06 C*07:02 | T98G | 6901 | 1876 |
| LNT-229+DAC | 27412690 | MULTIALLELIC-OLD | A*03:01 B*35:01 C*04:01 | LNT-229 | 7994 | 1477 |
| FM-82 | 27600516 | MULTIALLELIC-OLD | A*02:01 A*01:01 B*08:01<br>B*15:01 C*03:04 C*07:01 | MELANOMA_CELL_LINE | 3565 | 655 |
| FM-93/2 | 27600516 | MULTIALLELIC-OLD | A*02:01 A*26:01 B*40:01<br>B*44:02 C*03:04 C*05:01 | MELANOMA_CELL_LINE | 3289 | 899 |
| MEL-624 | 27600516 | MULTIALLELIC-OLD | A*02:01 A*03:01 B*07:02<br>B*14:01 C*07:02 C*08:02 | MELANOMA_CELL_LINE | 2402 | 373 |
| MEWO | 27600516 | MULTIALLELIC-OLD | A*02:01 A*26:01 B*14:02<br>B*38:01 C*08:02 C*12:03 | MEWO | 1679 | 495 |
| SK-MEL-5 | 27600516 | MULTIALLELIC-OLD | A*02:01 A*11:01 B*40:01<br>C*03:03 | SK-MEL-5 | 3028 | 598 |
| MEL-8 | 27869121 | MULTIALLELIC-OLD | A*01:01 A*03:01 B*07:02<br>B*08:01 C*07:01 C*07:02 | MELANOMA_MET | 6297 | 819 |
| MEL-12 | 27869121 | MULTIALLELIC-OLD | A*01:01 B*08:01 C*07:01 | MELANOMA_MET | 3776 | 959 |
| MEL-15 | 27869121 | MULTIALLELIC-OLD | A*03:01 A*68:01 B*27:05<br>B*35:03 C*02:02 C*04:01 | MELANOMA_MET | 21979 | 5841 |
| MEL-16 | 27869121 | MULTIALLELIC-OLD | A*01:01 A*24:02 B*07:02<br>B*08:01 C*07:01 C*07:02 | MELANOMA_MET | 12073 | 3113 |
| APHER1 | 28832583 | MULTIALLELIC-OLD | A*03:01 A*29:02 B*44:02<br>B*44:03 C*12:03 C*16:01 | LEUKAPHERESIS | 6163 | 1572 |
| APHER6 | 28832583 | MULTIALLELIC-OLD | A*02:01 A*03:01 B*07:02<br>C*07:02 | LEUKAPHERESIS | 1972 | 330 |
| CD165 | 28832583 | MULTIALLELIC-OLD | A*02:05 A*24:02 B*15:01<br>B*50:01 C*03:03 C*06:02 | B-CELL | 5381 | 1498 |
| CM467 | 28832583 | MULTIALLELIC-OLD | A*01:01 A*24:02 B*13:02<br>B*39:06 C*06:02 C*12:03 | B-CELL | 7408 | 3464 |
| GD149 | 28832583 | MULTIALLELIC-OLD | A*01:01 A*24:02 B*38:01<br>B*44:03 C*06:02 C*12:03 | B-CELL | 9760 | 3637 |
| MD155 | 28832583 | MULTIALLELIC-OLD | A*02:01 A*24:02 B*15:01<br>B*18:01 C*03:03 C*07:01 | B-CELL | 4393 | 768 |
| PD42 | 28832583 | MULTIALLELIC-OLD | A*02:06 A*24:02 B*07:02<br>B*55:01 C*01:02 C*07:02 | B-CELL | 2586 | 457 |
| RA957 | 28832583 | MULTIALLELIC-OLD | A*02:20 A*68:01 B*35:03<br>B*39:01 C*04:01 C*07:02 | B-CELL | 11052 | 5998 |
| TIL1 | 28832583 | MULTIALLELIC-OLD | A*02:01 A*02:01 B*18:01<br>B*38:01 C*05:01 | TIL | 5457 | 2070 |
| TIL3 | 28832583 | MULTIALLELIC-OLD | A*01:01 A*23:01 B*07:02<br>B*15:01 C*12:03 C*14:02 | TIL | 8804 | 2385 |
| PAT_AC2 | 28832583 | MULTIALLELIC-OLD | A*03:01 A*32:01 B*27:05<br>B*45:01 | B-LCL | 1388 | 124 |
| PAT_C | 28832583 | MULTIALLELIC-OLD | A*02:01 A*03:01 B*07:02<br>C*07:02 | B-LCL | 3182 | 287 |
| PAT_CELG | 28832583 | MULTIALLELIC-OLD | A*02:01 A*24:02 B*15:01<br>B*73:01 C*03:03 C*15:05 | B-LCL | 4143 | 942 |
| PAT_CP2 | 28832583 | MULTIALLELIC-OLD | A*11:01 B*14:02 B*44:02 | B-LCL | 1805 | 176 |
| PAT_FL | 28832583 | MULTIALLELIC-OLD | A*03:01 A*11:01 B*44:03<br>B*50:01 | B-LCL | 3768 | 539 |
| PAT_J | 28832583 | MULTIALLELIC-OLD | A*02:01 A*03:01 B*07:02<br>C*07:02 | B-LCL | 2696 | 256 |
| PAT_JPB3 | 28832583 | MULTIALLELIC-OLD | A*02:01 A*11:01 B*27:05<br>B*56:01 | B-LCL | 1994 | 129 |
| PAT_JT2 | 28832583 | MULTIALLELIC-OLD | A*11:01 B*18:03 B*35:01 | B-LCL | 1472 | 177 |
| PAT_M | 28832583 | MULTIALLELIC-OLD | A*03:01 A*29:02 B*08:01<br>B*44:03 C*07:01 C*16:01 | B-LCL | 2499 | 137 |
| PAT_MA | 28832583 | MULTIALLELIC-OLD | A*02:01 A*29:02 B*44:03<br>B*57:01 C*07:01 C*16:01 | B-LCL | 3869 | 489 |
| PAT_ML | 28832583 | MULTIALLELIC-OLD | A*02:01 A*11:01 B*40:01<br>B*44:03 | B-LCL | 3256 | 219 |
| PAT_NS2 | 28832583 | MULTIALLELIC-OLD | A*02:01 B*13:02 B*41:01 | B-LCL | 648 | 85 |
| PAT_NT | 28832583 | MULTIALLELIC-OLD | A*01:01 A*32:01 B*08:01 | B-LCL | 2306 | 349 |
| PAT_PF1 | 28832583 | MULTIALLELIC-OLD | A*01:01 A*02:01 B*07:02<br>B*44:03 C*07:02 C*16:01 | B-LCL | 4851 | 413 |
| PAT_R | 28832583 | MULTIALLELIC-OLD | A*03:01 A*29:02 B*08:01<br>B*44:03 C*07:01 C*16:01 | B-LCL | 2396 | 126 |
| PAT_RT | 28832583 | MULTIALLELIC-OLD | A*01:01 A*02:01 B*18:01<br>B*39:24 C*05:01 C*07:01 | B-LCL | 2722 | 700 |
| PAT_SR | 28832583 | MULTIALLELIC-OLD | A*02:01 A*23:01 B*18:01<br>B*44:03 | B-LCL | 2770 | 282 |
| PAT_ST | 28832583 | MULTIALLELIC-OLD | A*03:01 A*24:02 B*07:02<br>B*27:05 | B-LCL | 1263 | 40 |
| 11-002-S1-TISSUE | 31154438 | MULTIALLELIC-RECEN<br>T | A*03:01 A*24:02 B*35:03<br>B*44:02 C*12:03 C*12:03 | GLIOBLASTOMA_TISSUE | 3303 | 969 |
| 10-002-S1-TISSUE | 31154438 | MULTIALLELIC-RECEN<br>T | A*02:01 A*31:01 B*13:02<br>B*58:01 C*06:02 C*07:01 | GLIOBLASTOMA_TISSUE | 980 | 431 |
| BCN-018-TISSUE | 31154438 | MULTIALLELIC-RECEN<br>T | A*02:01 A*29:01 B*07:02<br>B*27:05 C*01:02 C*15:05 | GLIOBLASTOMA_TISSUE | 2556 | 935 |
| CPH-09-TISSUE | 31154438 | MULTIALLELIC-RECEN<br>T | A*02:01 A*32:01 B*27:05<br>B*44:02 C*05:01 C*02:02 | GLIOBLASTOMA_TISSUE | 5281 | 1527 |
| CPH-07-TISSUE | 31154438 | MULTIALLELIC-RECEN<br>T | A*02:01 A*02:01 B*35:01<br>B*27:05 C*02:02 C*04:01 | GLIOBLASTOMA_TISSUE | 5589 | 1816 |
| 29/14-TISSUE | 31154438 | MULTIALLELIC-RECEN<br>T | A*02:01 A*32:01 B*40:01<br>B*13:02 C*03:04 C*06:02 | GLIOBLASTOMA_TISSUE | 5235 | 4049 |
| 637/13-TISSUE | 31154438 | MULTIALLELIC-RECEN<br>T | A*01:01 A*24:02 B*51:01<br>B*08:01 C*07:01 C*01:02 | GLIOBLASTOMA_TISSUE | 3345 | 2399 |
| LEIDEN-005-TISSUE | 31154438 | MULTIALLELIC-RECEN<br>T | A*02:01 A*25:01 B*35:01<br>B*18:01 C*12:03 C*04:01 | GLIOBLASTOMA_TISSUE | 6226 | 2330 |
| CPH-08-TISSUE | 31154438 | MULTIALLELIC-RECEN<br>T | A*32:01 A*26:01 B*38:01<br>B*40:02 C*02:02 C*12:03 | GLIOBLASTOMA_TISSUE | 6679 | 3036 |
| LEIDEN-004-TISSUE | 31154438 | MULTIALLELIC-RECEN<br>T | A*03:01 A*02:01 B*07:02<br>B*07:02 C*12:03 C*07:02 | GLIOBLASTOMA_TISSUE | 4705 | 1329 |
| KESKIN_DFCI-5283 | 31844290 | MULTIALLELIC-RECEN<br>T | A*01:01 A*02:01 B*07:02<br>B*08:01 C*07:01 C*07:02 | B-CELL | 5866 | 958 |
| KESKIN_DFCI-5328 | 31844290 | MULTIALLELIC-RECEN<br>T | A*02:06 A*24:02 B*08:01<br>B*51:01 C*07:02 C*14:02 | B-CELL | 308 | 50 |
| KESKIN_DFCI-5341 | 31844290 | MULTIALLELIC-RECEN<br>T | A*03:01 A*31:01 B*14:02<br>B*35:02 C*04:01 C*08:02 | B-CELL | 442 | 119 |
| KESKIN_CP-594_V1 | 31844290 | MULTIALLELIC-RECEN<br>T | A*02:01 A*24:02 B*35:03<br>B*44:02 C*05:01 C*12:03 | OVARY | 2581 | 478 |
| KESKIN_13240-015 | 31844290 | MULTIALLELIC-RECEN<br>T | A*02:01 A*02:02 B*13:02<br>B*40:02 C*02:02 C*06:02 | MELANOMA | 5170 | 1107 |
| KESKIN_13240-002 | 31844290 | MULTIALLELIC-RECEN<br>T | A*02:01 A*24:02 B*15:01<br>B*44:02 C*05:01 C*07:02 | MELANOMA | 967 | 122 |
| KESKIN_13240-005 | 31844290 | MULTIALLELIC-RECEN<br>T | A*01:01 A*01:01 B*38:01<br>B*56:01 C*01:02 C*06:02 | MELANOMA | 3404 | 762 |
| KESKIN_13240-006 | 31844290 | MULTIALLELIC-RECEN<br>T | A*02:01 A*03:01 B*27:05<br>B*47:01 C*01:02 C*06:02 | MELANOMA | 2649 | 472 |
| KESKIN_H4198_BT187 | 31844290 | MULTIALLELIC-RECEN<br>T | A*01:01 A*32:01 B*15:01<br>B*15:01 C*03:03 C*03:04 | GLIOBLASTOMA_TISSUE | 6151 | 1444 |
| KESKIN_H4512_BT145 | 31844290 | MULTIALLELIC-RECEN<br>T | A*01:01 A*32:01 B*13:02<br>B*40:02 C*02:02 C*06:02 | GLIOBLASTOMA_TISSUE | 6140 | 1345 |

**Table S2.**
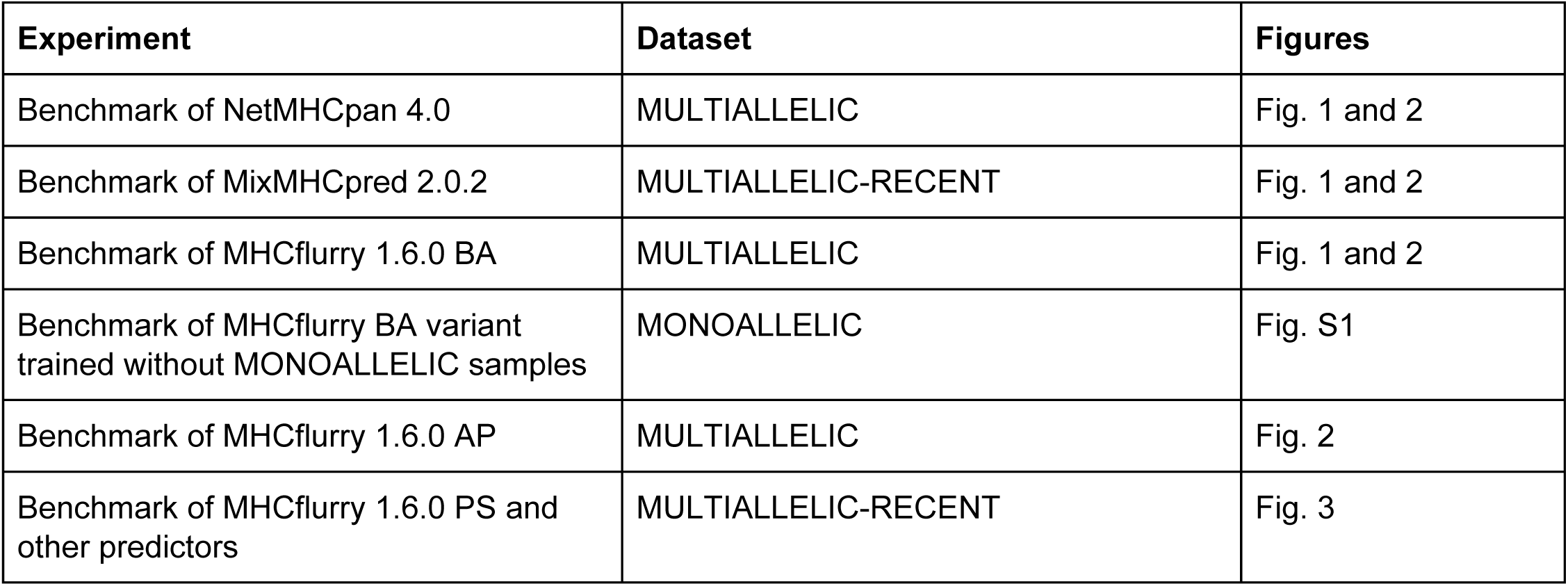
Summary of sample groups used for each benchmark experiment. The MULTIALLELIC set is the union of the MULTIALLELIC-OLD and MULTIALLELIC-RECENT samples.

**Table S3.**
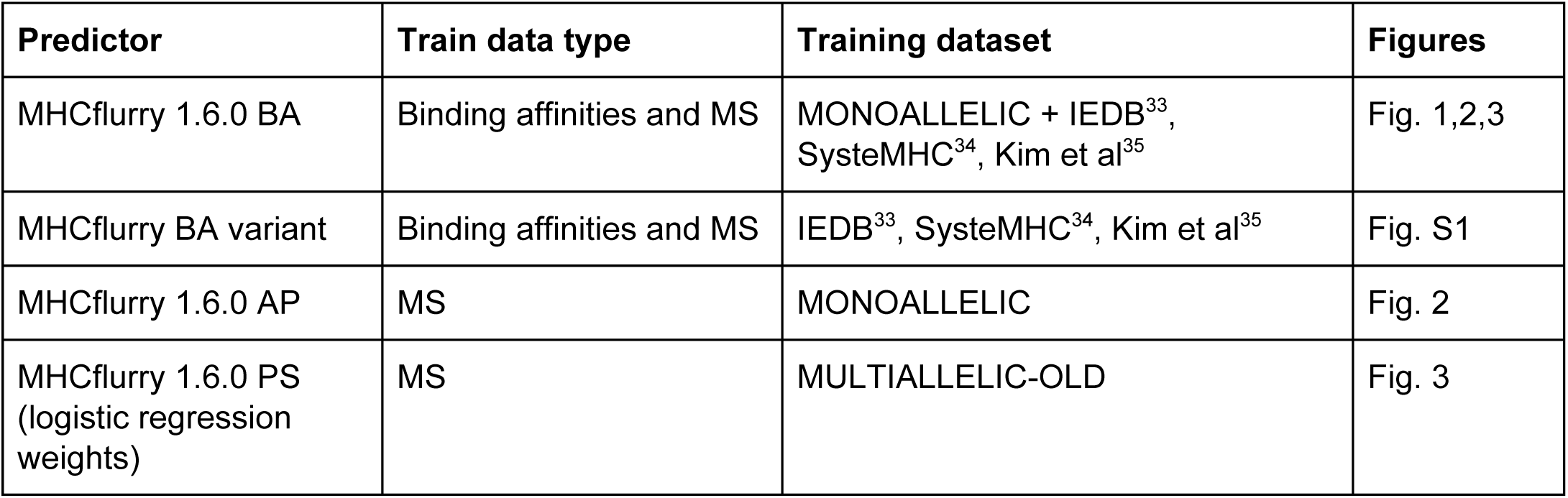
Summary of training datasets for the predictors developed in this work.

| Predictor | Train data type | Training dataset | Figures |
| --- | --- | --- | --- |
| MHCflurry 1.6.0 BA | Binding affinities and MS | MONOALLELIC + IEDB<sup>33</sup>,<br>SysteMHC<sup>34</sup>, Kim et al<sup>35</sup> | Fig. 1,2,3 |
| MHCflurry BA variant | Binding affinities and MS | IEDB<sup>33</sup>, SysteMHC<sup>34</sup>, Kim et al<sup>35</sup> | Fig. S1 |
| MHCflurry 1.6.0 AP | MS | MONOALLELIC | Fig. 2 |
| MHCflurry 1.6.0 PS<br>(logistic regression<br>weights) | MS | MULTIALLELIC-OLD | Fig. 3 |

**Fig S1:**
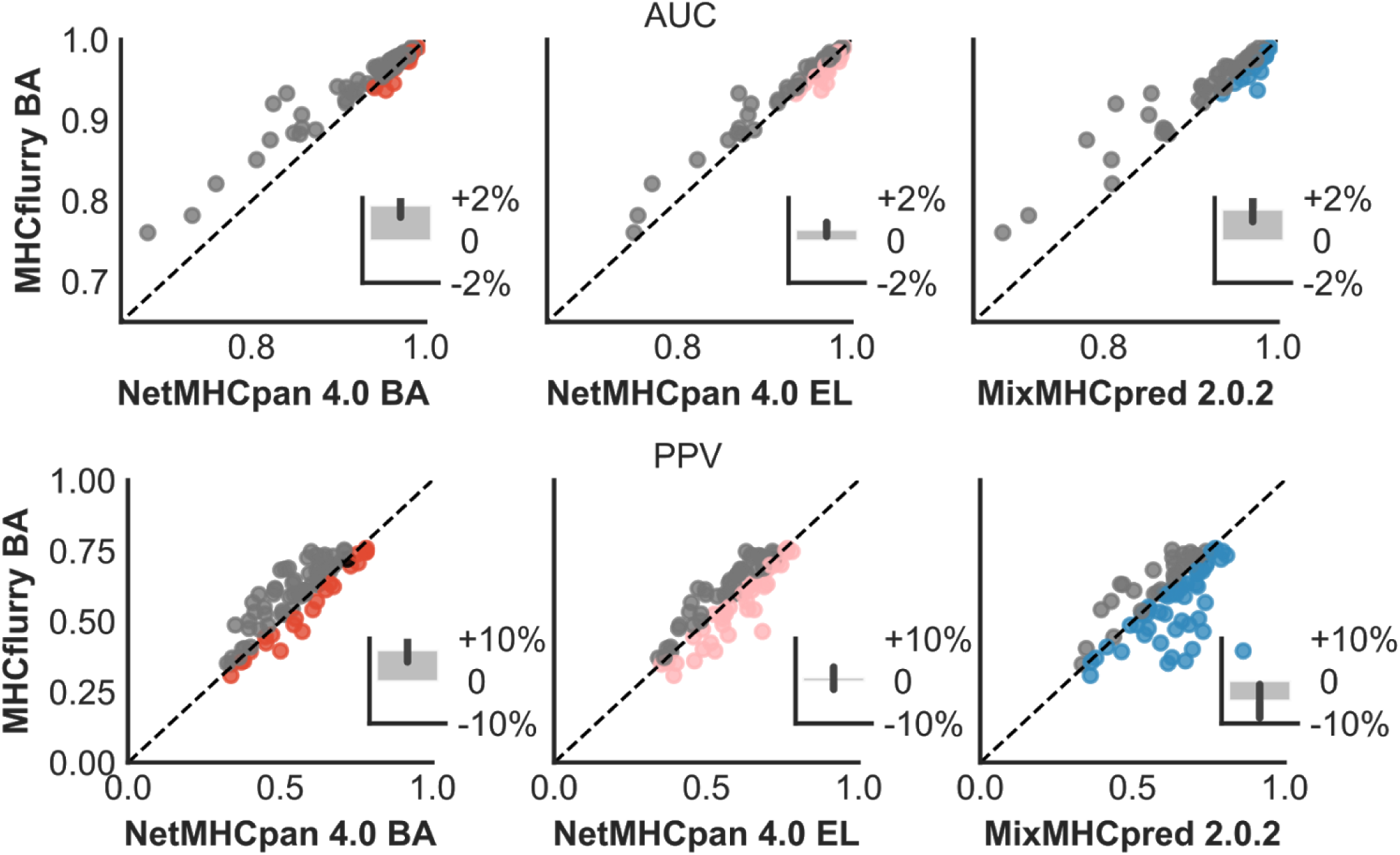
Comparison of MHCflurry BA and other predictors on the MONOALLELIC benchmark. A variant of MHCflurry BA trained without the MONOALLELIC benchmark datasets was used. The top row shows AUC; the bottom row shows PPV. The insets show the mean percent differences, with positive values indicating higher performance by MHCflurry BA. Error bars indicate 95% confidence intervals of the mean.

## Supplemental notes

### Supplemental Note 1

The approach we describe to train the AP predictor requires that the BA predictor does not fully model the antigen processing signals available in its MS training datasets. While the BA predictor likely includes a contribution from antigen processing, we note that several technical choices may have blunted its ability to learn these effects. The BA predictor’s training set includes 69% mass spec-identified ligands and 31% peptide-MHC I affinity measurements. The predictor therefore models a compromise between antigen processing-sensitive (MS ligand) and -insensitive (affinity measurement) training data. The influence of MS data is expected to be further diluted at the extreme strong-binder segment of the BA predictor’s output because MS hits are assigned an ic_50_ of “< 50 nM” in the training set, i.e. any assigned affinity tighter than 50 nM results in zero contribution to the training loss. This means that MS training data does not guide the relative ranking (exact ic_50_) of strong binders, potentially the regime where antigen processing signals may make the greatest difference. This effect likely also plays a role in the relatively lackluster performance of the BA predictor when assessed by the PPV metric -- which emphasizes the rank-order of peptides at the extreme high-end of the predictor’s output -- despite good performance on AUC. Finally, an important difference between the BA and AP predictors is that the AP predictor uses peptides from the proteome as decoys (unobserved non-binders), whereas the BA predictor uses random sequences sampled according to the same amino acid distribution as the hits. The AP predictor strategy is expected to be more realistic and informative, at the cost of giving the AP predictor more opportunity to learn MS biases.

## Supplemental data

These files have been deposited in Mendeley Data at https://data.mendeley.com/datasets/zx3kjzc3yx.

**Additional File 1.** MULTIALLELIC benchmark dataset with predictions (CSV).

**Additional File 2.** MONOALLELIC benchmark dataset with predictions (CSV).

**Additional File 3.** Training data for MHCflurry 1.6.0 binding affinity (BA) predictors (CSV).

**Additional File 4.** Training data for the variant of MHCflurry BA evaluated on the MONOALLELIC benchmark in Fig. S1 (CSV).

**Additional File 5.** Training data for MHCflurry 1.6.0 antigen processing (AP) predictors (CSV).

**Additional File 6.** Training data for MHCflurry 1.6.0 presentation score (PS) predictors (CSV).

**Additional File 7.** Position weight matrices for the MHCflurry antigen processing predictors (XLSX).

**Additional File 8.** Antigen processing predictions for proteosome-cleaved peptides identified using “mass spectrometry analysis of proteolytic peptides” (MAPP) by Wolf-Levy et al^15^.

